# Ecological specialisation, rather than the island effect, explains morphological diversification in an ancient radiation of geckos

**DOI:** 10.1101/2021.07.30.454424

**Authors:** Héctor Tejero-Cicuéndez, Marc Simó-Riudalbas, Iris Menéndez, Salvador Carranza

## Abstract

Island colonists are often assumed to experience higher levels of phenotypic diversification than continental taxa. However, empirical evidence has uncovered exceptions to this “island effect”. Here, we tested this pattern using the geckos of the genus *Pristurus* from continental Arabia and Africa and the Socotra Archipelago. Using a recently published phylogeny and an extensive morphological dataset, we explore the differences in phenotypic evolution between Socotran and continental taxa. Moreover, we reconstructed ancestral habitat occupancy to examine if ecological specialisation is correlated with morphological change, comparing phenotypic disparity and trait evolution between habitats. We found a heterogeneous outcome of island colonisation. Namely, only one of the three colonisation events resulted in a body size increase. However, in general, Socotran species do not present higher levels or rates of morphological diversification than continental groups. Instead, habitat specialisation explains better the body size and shape evolution in *Pristurus*. Particularly, the colonisation of ground habitats appears as the main driver of morphological change, producing the highest disparity and evolutionary rates. Additionally, arboreal species show very similar body size and head proportions. These results reveal a determinant role of ecological mechanisms in morphological evolution and corroborate the complexity of ecomorphological dynamics in continent-island systems.

## INTRODUCTION

The life history and population biology of continental and insular taxa of a specific evolutionary radiation are fundamentally distinct [1,2]. In the continent, communities are often assumed to be complex and composed of many species that share a long history of coevolution [3]. In such a scenario, most of the ecological niches will be filled, and high levels of interspecific competition are expected [4]. These factors, together with higher predation pressures, will tend to limit niche expansion and, consequently, morphological diversification [5]. In contrast, insular groups are usually exposed to higher levels of ecological opportunity and thus, they can occupy the new or relatively unexploited adaptive landscapes that islands provide [6,7]. As a result, island species may experience increased rates of phenotypic diversification and higher levels of morphological disparity compared to continental taxa [8]. Moreover, the divergence in body size found in insular taxa relative to their continental counterparts is a pervasive and widely studied pattern across vertebrates, known as the “island rule” [9]. However, empirical evidence outlines a more complex scenario in which island colonists might not necessarily experience great levels of evolutionary divergence [10]. Instead, these potential outcomes of island colonisation –– i.e., body size and shape divergence, increased phenotypic disparity, and increased rates of morphological evolution; from now on referred to as “island effect” –– might depend on multiple *extrinsic* factors (mostly modulated by the geography or geology of the island), as well as *intrinsic* factors (i.e., the biological characteristics of the group concerned) [3,11]. Thus, ecological specialisation is expected to be more pronounced when island colonisation results in an expansion into novel ecological contexts, and such specialisation might carry morphological changes depending on species and system-specific factors [4,6].

Whether ecological specialisation follows island colonisation or not, the study of habitat occupancy is essential to understand the evolution of associated traits and the structuring of ecological communities [12]. In particular, specialisation in substrate use can promote morphological diversity through deterministic body size evolution and diversification [13]. Moreover, microhabitat use can be strongly correlated with convergent phenotypic evolution resulting in recurrent ecomorphs beyond the effect of history and clade membership [14].

Arid regions, generally considered relatively depauperate in terms of animal diversity, have been successfully inhabited by some vertebrate groups and harbour especially high levels of reptile diversity. Among reptiles, geckos are particularly prominent due to their outstanding diversity in ecological features, exhibiting a wide variety of morphological and behavioural adaptations [15]. Afro-Arabian geckos have been recently prominent in studies of the role of arid biomes in generating biodiversity [16–21]. However, the outcomes of morphological diversification in these animals have only been properly investigated within the genus *Hemidactylus*, which is the best-studied Arabian reptile group with well-resolved taxonomy and reliably reconstructed biogeographic history [18,19,22–27]. In particular, two recent studies including continental and insular taxa from the Socotra Archipelago indicated that the genus *Hemidactylus* shows an island effect at least regarding body size evolution [28,29]. In contrast, the geckos of the genus *Pristurus*, which have also colonised and diversified within the same archipelago, seem to show lower rates of body size diversification in Socotra than other genera in the archipelago, but also compared to their continental relatives [28]. Despite these preliminary results, a more nuanced analysis of the morphological evolution in *Pristurus*, including undescribed diversity, morphological and ecological data, is still lacking. Interestingly, the genus *Pristurus* not only has representatives in both the Socotra Archipelago and continental Africa and Arabia, but it also has diversified ecologically to exploit a variety of habitats, including rocky and sandy surfaces, gravel plains, and trees [30,31].

Here we use a recently inferred phylogenetic assessment of Afro-Arabian reptiles including undescribed diversity, together with an extensive morphological sampling and detailed ecological information, to explore the morphological evolution in *Pristurus* geckos. Specifically, we test alternative scenarios of body size and shape evolution in this genus, to determine the role of the colonisation of the Socotra Archipelago and the ecological specialisation in generating the morphological diversity observed. The independent diversification of both Socotran and continental taxa, the ecological and behavioural diversity, and the unique phenotypic dataset compiled in this study, make this group of geckos an exceptional system to investigate keystone dynamics in evolutionary biology such as the island effect and ecological adaptation, and their impact on morphological evolution.

## MATERIALS AND METHODS

### Phylogenetic and ecological data

We used a recently published phylogenetic tree of Afro-Arabian squamates [32]. This tree contains all the species of *Pristurus* for which there exists genetic data, including some species currently in the process of being described, resulting in a total of 30 species. We extracted the *Pristurus* clade from the squamate tree, both for the consensus and for 1,000 trees randomly selected from the posterior distribution generated in the cited study. Using a sample of posterior trees allowed us to take into account the phylogenetic and temporal uncertainty in the subsequent analyses.

Each species was defined as present in the Socotra Archipelago or in the continent (Africa and Arabia). For habitat specialisation, each species was categorized based on substrate preferences into one of three groups: ground-dwelling, rock climber, or arboreal [30,33]. Additionally, the ground-dwelling species were divided into “soft-ground” (sandy surfaces) and “hard-ground” (gravel plains) categories to further characterize the morphological adaptations to each type of ground habitat. Nevertheless, the disparity dynamics and rates of trait evolution were estimated with the original categorizations (continent -Socotra; and the three habitat states) which, due to the limited number of species, were more appropriate for the analyses.

### Ancestral reconstructions

We studied the colonisation of the Socotra Archipelago and the habitat specialisation through time by reconstructing ancestral states across the phylogeny. First, we fitted several models of character evolution across the phylogeny in order to select the best-fit model for both traits. With such a purpose, we used the function *fitDiscrete* from the R package geiger v2.0 [34,35]. We fitted three models: an equal-rates model (ER), a symmetrical model (SYM), and an all-rates-different model (ARD). We selected the best-fit model in each case based on the Akaike information criterion, correcting for small sample size (AICc; Akaike 1973). We then used the function *make*.*simmap* from the R package phytools [37], which simulates plausible stochastic character histories after fitting a continuous-time reversible Markov model for the evolution of the character states assigned to the tips of the tree. We ran 1,000 simulations with the previously selected model of character evolution (ER model for both traits). Additionally, we randomly selected 100 trees from the posterior distribution, and we ran 100 stochastic character histories on each of them for both traits (presence in Socotra and habitat occupancy).

### Phenotypic data

For the 30 species included in the phylogenetic tree, a total of 697 specimens were examined and measured, with a minimum of one, a maximum of 56, and a mean of 23 specimens per species (Table S1). All vouchers were obtained from the following collections: Institute of Evolutionary Biology (CSIC-UPF), Barcelona, Spain (IBE), Natural History Museum, London, UK (BM), Museo Civico di Storia Naturale, Carmagnola, Turin, Italy (MCCI), Università di Firenze, Museo Zoologico “La Specola”, Firenze, Italy (MZUF), Oman Natural History Museum (ONHM), Laboratoire de Biogéographie et Écologie des Vertébrés de l’École Pratique des Hautes Etudes, Montpellier, France (BEV), and National Museum Prague, Czech Republic (NMP). The following measurements were taken by the same person (MSR) using a digital caliper with accuracy to the nearest 0.1 mm: snout-vent length (SVL; distance from the tip of the snout to the cloaca), trunk length (TrL; distance between forelimb and hindlimb measured posterior to the forelimb and anterior to the hindlimb insertion points), head length (HL; taken axially from tip of the snout to the anterior ear border), head width (HW; taken at anterior ear border), head height (HH; taken laterally at anterior ear border), humerus length (Lhu; from elbow to axilla), ulna length (Lun; from wrist to elbow), femur length (Lfe; from knee to groin) and tibia length (Ltb; from ankle to knee). Tail length was not measured because most of the specimens had regenerated tails or had lost it.

### Morphological differentiation

As body size and shape evolution might be affected by the colonisation of Socotra and/or the ecological specialisation, we investigated whether Socotran colonists are morphologically distinct from their relatives in the continent (one potential outcome under the “island effect” framework) and likewise whether the differential habitat use is reflected in species morphology. For each species, the mean of each morphological variable was calculated and log_10_-transformed in order to improve normality and homoscedasticity prior to subsequent analyses.

To characterize shape differentiation, we performed a phylogenetic regression of each trait on snout-vent length (SVL) to remove the effect of the body size on the other variables. The residuals of these regressions were used to implement a phylogenetically controlled principal component analysis (pPCA) using the functions *phyl*.*resid* and *phyl*.*pca* from the R package phytools with the method set to ‘lambda’ [37]. We used the principal components representing 75% of the cumulative proportion of variance as shape variables and we visualised the 2D shape morphospace. Additionally, we performed a principal component analysis (PCA) with the shape data from all the specimens measured, after correcting for body size through regressions on SVL similarly to our processing of the species data. We generated per-species boxplots of size and shape variation with the specimen data.

For body size, we used the function *phenogram* from phytools [37] to map and visualize variation in SVL across the species tree. For all the visualisations, we categorised the species according to their presence in the Socotra Archipelago (Socotra or continent) and to their habitat use (ground, rock or tree) separately, to have a detailed perspective of the extent of the morphological differentiation undergone in each category. Furthermore, to test the island rule prediction that body size in insular species is different from the continent, we performed a phylogenetic ANOVA with the *phylANOVA* function from phytools [37]. Additionally, according to the island rule, we would expect to see body size differences before and after a colonisation event. To test this, we performed paired t-tests on the estimated body size for each pair of parental and descendent nodes in which a change from continent to Socotra occurred. We used the function *OUwie*.*anc* from the R package OUwie [38] to estimate the ancestral body size under a multi-rate Brownian Motion model, which was the best-fit model for body size evolution (see below “Differences in tempos of phenotypic diversification”).

### Exploring differences in phenotypic disparities

Since one of the main possible outcomes of island colonisation and/or ecological specialisation is the increase in phenotypic disparity, we tested this assumption following previous research on other geckos [29] and defining disparity as the average squared Euclidean distance between all pairs of species in a group for a given continuous variable [7], in our case body size (SVL) and two variables of shape (pPC1 and pPC2). First, we tested whether disparity is higher in Socotran species than in the continent, which would indicate the existence of an island effect. Then, considering our results of morphological differentiation according to habitat use, in which ground-dwelling species are widespread throughout most of the morphospace of the genus, we tested whether the phenotype of ground-dwelling species is significantly more disparate than that of species in all the other habitats together (i.e., rock and arboreal species). We calculated the observed disparity ratios (Socotra/continent and ground/no-ground) for each morphological variable, using the function *disparity* from the R package geiger [34]. In the case of higher disparity in Socotra or in ground species (disparity ratio Socotra/continent or ground/no-ground higher than 1), we then compared the observed ratios with a null distribution of disparity ratios obtained from 10,000 simulations of the evolution of a continuous character according to a Brownian motion model across the phylogeny. These simulations were performed by applying the function *sim*.*char* after estimating the empirical rate parameter for body size and shape from the best-fit model (Brownian motion) with the function *fitContinuous*, both from the package geiger [34]. This approach allowed us to test, in the case of an observed higher disparity in Socotran or ground species, whether this is a significant increase considering the rate of evolution of the character, or rather this is not evidence of effectively increased disparity. In order to test the sensitivity of the results to low intraspecific sampling, we performed the analyses described above also with a dataset in which the species with less than five specimens had been removed: *Pristurus adrarensis, P. flavipunctatus, P*. sp. 4, *P*. sp. 9, and *P*. sp. 12 (see Table S1).

### Differences in tempos of phenotypic diversification

In order to test the effect of Socotran colonisation and ecological specialisation in the tempo of phenotypic evolution, we fitted different models of character evolution across the phylogeny in which the evolutionary rates of body size and shape might or might not differ between categories (i.e., between Socotra and continent, and between ground, rock and tree habitats). For body size, we used the R package OUwie [38] to fit three alternative models: BM1 (Brownian motion single rate, i.e., assuming one single rate regime for all lineages in the phylogeny), OU1 (Ornstein-Uhlenbeck single-rate value with a phenotypic optimum and a selective pressure towards it), and BMS (Brownian motion multi-rate model, with different rate values for each of the regimes specified, i.e., Socotran different from continental lineages, and differences between habitats). Similarly, we studied the rates of phenotypic evolution for body shape, but in this case we fitted multivariate models including the first two principal components resulting from the phylogenetic PCA (pPC1 and pPC2; 77% of the variance, see Results section). We used the R package mvMorph [39] to fit four alternative models: BM1, OU1, and BMM (analogous to BMS in the OUwie package), and OUM (multi-rate Ornstein-Uhlenbeck model). We fitted these models in the 1,000 stochastic character maps generated for the consensus tree, and also in the 100 character maps on each of the 100 posterior trees used to reconstruct ancestral states (see above Ancestral reconstructions). We then selected the best-fit models based on the AICc distributions and means, and we extracted the distributions of rate values estimated for each regime (Socotra, continent, ground, rock and tree) in the multi-rate models, to unravel the effect of each trait in morphological rates. All the visualisations of the disparity and phenotypic rate analyses were built with the R packages ggplot2 [40], patchwork [41], and cowplot [42].

## RESULTS

### Ancestral reconstructions and morphological variation

The ancestral reconstructions following an equal-rates model (ER), with the probabilities of each state in ancestral nodes (Socotra and continent; ground, rock, and tree) can be found in the Supplementary Material (Fig. S1). For the Socotra-continent reconstruction, the most probable ancestral state for the root was “continent” (continent: 0.649; Socotra: 0.351). The ecological reconstruction shows rocky habitats as the ancestral state in *Pristurus* (rock: 0.978; ground: 0.012; tree: 0.010), with several transitions to arboreal habitats and one colonisation of the ground in the ancestor of the clade known as “Spatalura group” (Arnold 1993, 2009; Fig. S1). One of the subclades of this group later colonised more compact, harder substrates, shown in our more detailed analysis separating soft- and hard-ground species (Fig. S1).

The pPCA analysis of body shape resulted in two first components explaining 77% of the total variance: pPC1 (61% of the variance) mainly representing limb dimensions (variables Lhu, Lun, Lfe, Ltb), and pPC2 (16% of the variance) mostly representing head proportions (variables HL, HW, HH) (Fig. 1a; Table S2; see Fig. S2 and Table S3 for results on body size and shape differentiation using the specimen data and PCA). The morphospace occupied by continental species is notably larger than that occupied by Socotran species, and they overlap almost completely (Fig. 1a left). When visualizing the shape phylomorphospace along with habitat categories, we observe a wide portion occupied by the ground-dwelling species, especially in pPC2 (head dimensions) (Fig. 1a right). These eight species of the “Spatalura group” essentially occupy almost as much of the morphospace as all the rest of the species together, with morphologies specialised to arboreal habits localized in a narrow area, especially for head proportions (Fig. 1a right).

**Figure 1.**
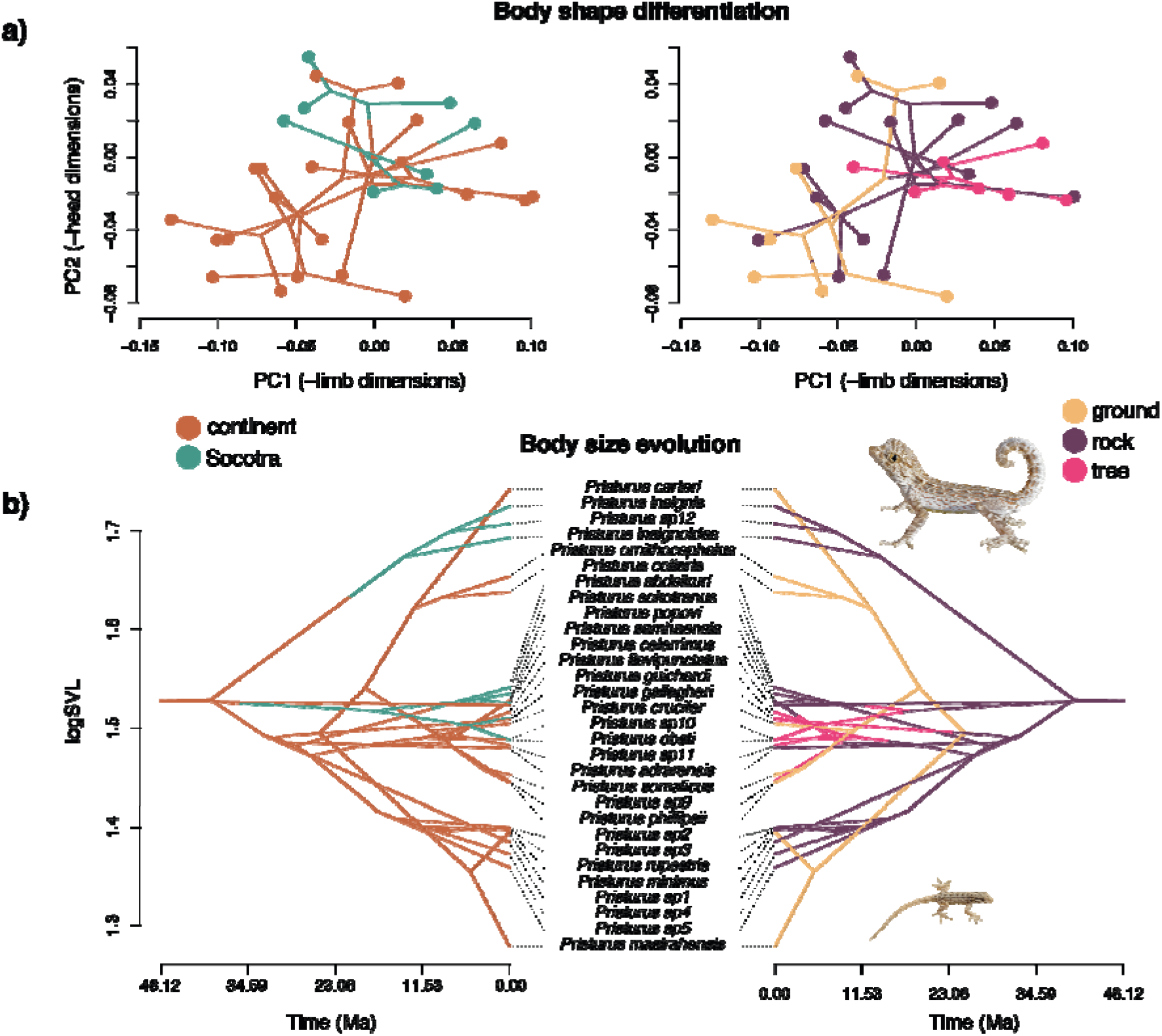
Morphological variability in *Pristurus*, with insight from insularity (left) and habitat use (right). a) Morphospace with phylogenetic relationship between the species, showing body shape differentiation. b) Traitgram showing body size (SVL) through time on the summary phylogenetic tree of *Pristurus*, mapped by the discrete categories of presence in Socotra or the continent (left) and by ecological specialisation (right). Photos (proportional to species’ SVL): *Pristurus carteri* (top) and *P. masirahensis* (bottom), the largest and smallest species of the genus, respectively.

We found a similar pattern for body size. Body size variability of Socotran species is completely contained in the range occupied by continental species (Fig. 1b left). On the contrary, ground-dwelling species show a size variability higher than all the species from other habitats together, being the largest and the smallest species of *Pristurus* specialised to ground habitats (Fig. 1b right). As with head proportions, arboreal species have apparently constrained body sizes, being restricted to specific intermediate values within the genus’ size range. The phylogenetic ANOVA showed no statistical differences in body size between Socotran and continental species (F = 7.535, p = 0.215). Although one of the colonisation events of the Socotra Archipelago was followed by an apparent increase in body size (the clade composed by *Pristurus insignis, P. insignoides*, and *P*. sp. 12; see Fig. S3), our paired t-test analysis did not find significant differences between parental and descendant nodes in which a change from continent to Socotra occurred (t = -1.201, p = 0.353; Fig. S3).

When separating ground species into hard- and soft-ground habitats, we observed a clear morphological differentiation, especially in body size. Hard-ground species seem to be highly specialised, with some of the largest body sizes of the genus, long limbs and large heads (Fig. S4).

### Phenotypic disparity

We found that the morphological disparity in the Socotra Archipelago is lower than in the continent for the three variables, with disparity ratios Socotra/continent below 1 (SVL: 0.88; pPC1: 0.52; pPC2: 0.53). When comparing disparity between ground and no-ground habitats, we found a higher observed disparity in ground for body size (SVL) and head proportions (pPC2), with disparity ratios ground/no-ground of 2.25 for SVL, 0.86 for pPC1, and 2.47 for pPC2. Furthermore, both for size and head proportions, the increased disparity in ground habitats was significant compared with the null distribution of simulated disparity ratios (p_size_ = 0.03; p_head_ = 0.01; Fig. 2). These results were virtually identical to those from the analyses performed after removing the species with less than five specimens (Table S4).

**Figure 2.**
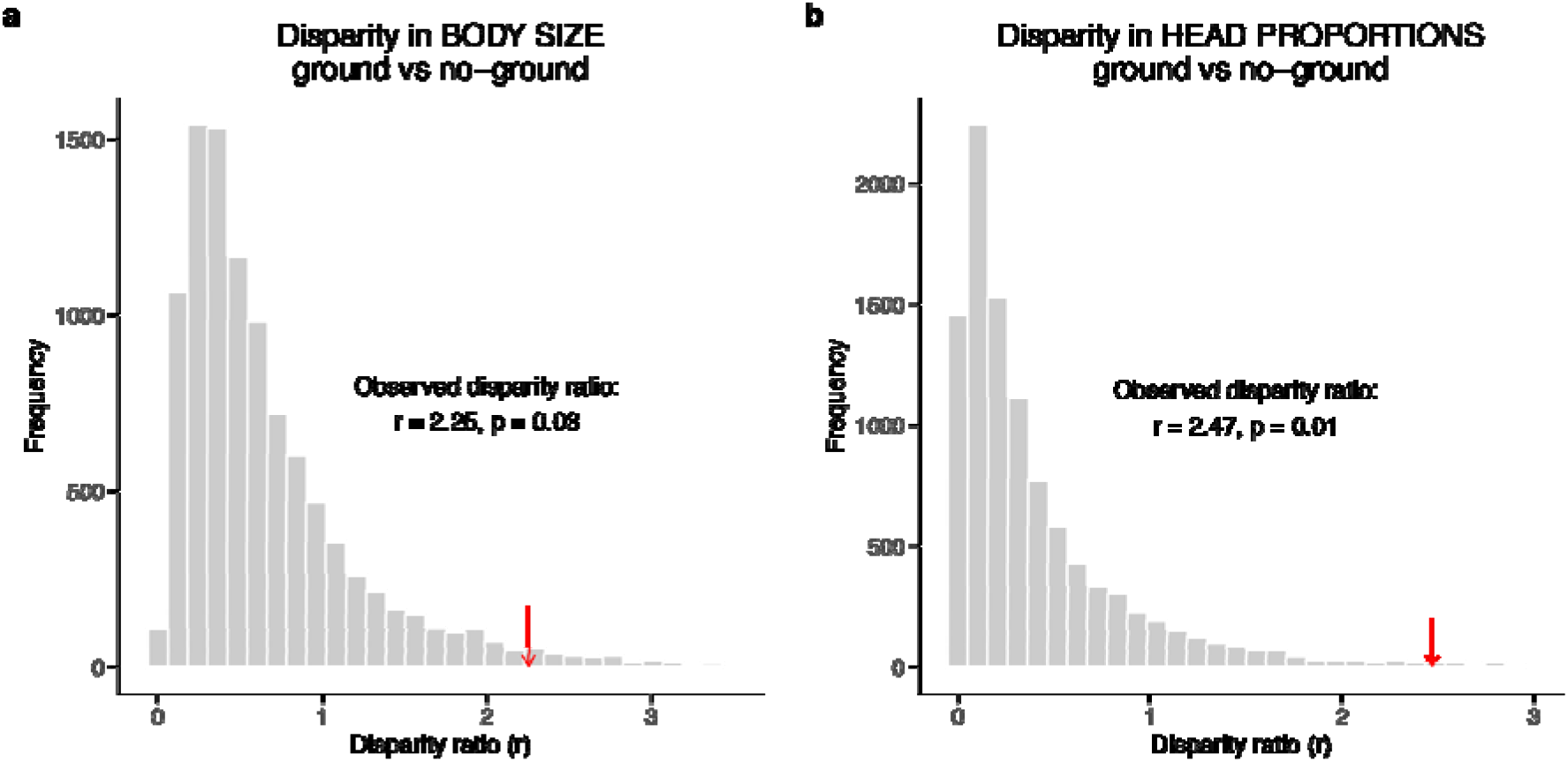
Observed (red arrows) and simulated (grey bars) ratios of phenotypic disparity between ground versus no-ground habitats. a) Body size disparity ratios. b) Head proportions (pPC2) disparity ratios.

### Rates of morphological evolution

For body size, a multi-rate Brownian motion model (BMS) was the best fit both for presence in Socotra and for ecological specialisation (lowest AICc; Fig. 3a), suggesting differences in the rate of morphological evolution across discrete categories (i.e., Socotra vs. continent, and different habitats). For body shape, however, we did not find evidence for differences in evolutionary rates, being the single-rate Brownian motion model (BM1) the best fit both for limb (pPC1) and head (pPC2) dimensions, although the overlap across all models was considerable (Fig. 3b).

**Figure 3.**
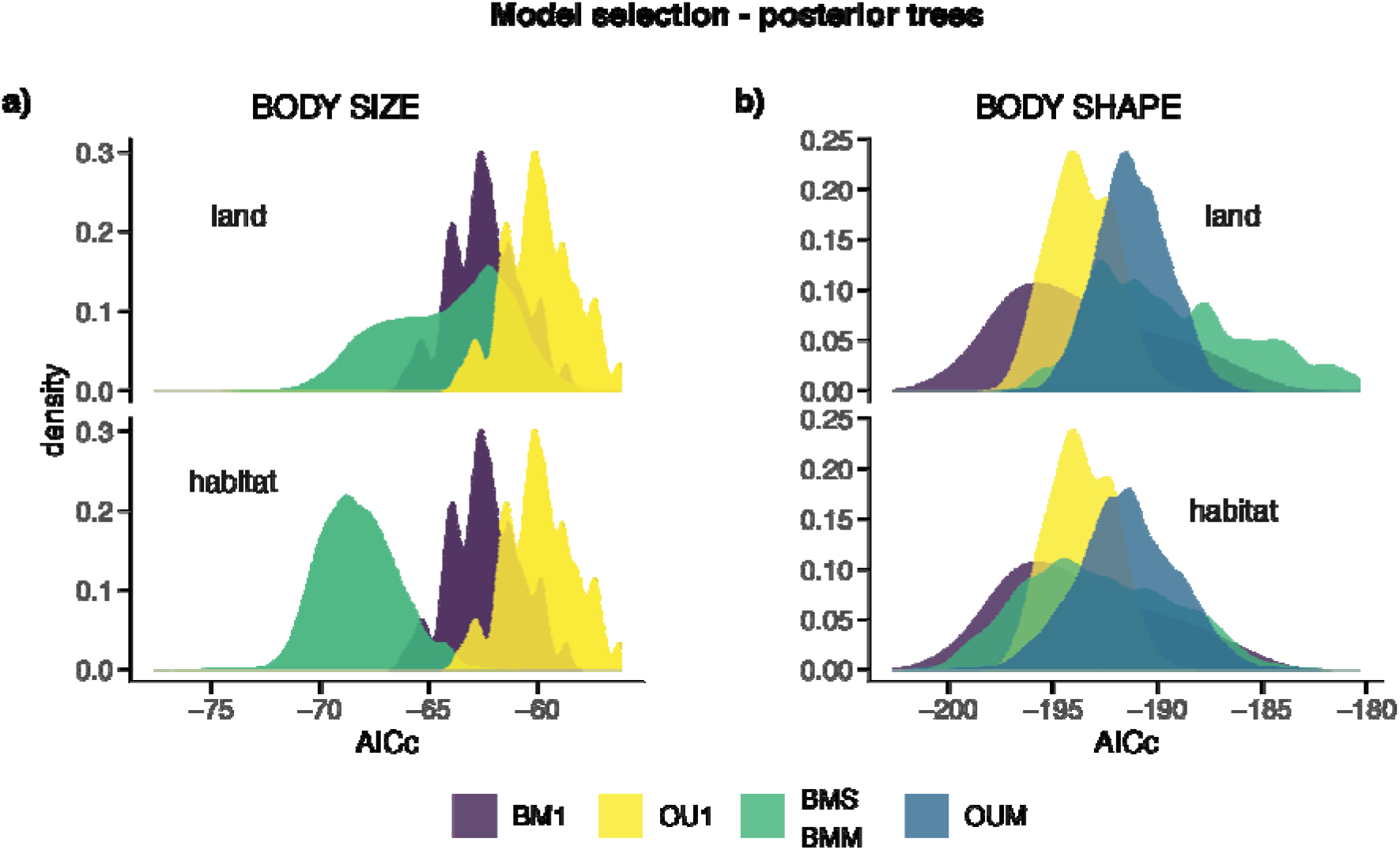
AICc distributions from the model fitting for a) body size and b) shape evolution of the genus *Pristurus*. These results correspond to model fitting on 100 stochastic character maps (insularity in the top panels and habitat use in the bottom) on 100 trees from the phylogenetic posterior distribution. BM1: Brownian motion single rate. OU1: Ornstein-Uhlenbeck single rate. BMS (OUwie) / BMM (mvMorph): Brownian motion multi-rate. OUM: Ornstein-Uhlenbeck multi-rate. For body size, the best supported model is a Brownian motion with rate heterogeneity across categories. For body shape, a single-rate Brownian motion model was the best-fit, although there is an extensive overlap across all models.

We extracted the rates of body size evolution from the Brownian motion multi-rate models, and we found that Socotran species present lower rates than species in the continent (Fig. 4a top), although the bimodal distribution of the rates in Socotran taxa suggests rate heterogeneity among those clades. For ecological specialisation, we found increased rates of body size evolution in the ground-dwelling species relative to the other habitats, with arboreal habitats showing the lowest rates (Fig. 4a bottom). We also extracted per-category rates of body shape evolution according to the BMM model in mvMorph, even though the multi-rate models were not the best supported, and we found a similar scenario, in which shape evolution (both for limbs and for head proportions) would be notably faster in ground-dwelling species (Fig. 4b bottom). Results from the analyses performed with the 1,000 stochastic character maps on the consensus tree and with the 100 maps on each of 100 posterior trees lead to the same conclusions, so we show the ones from the posterior trees on the main text. Results from analyses with the consensus tree can be found in the Supplementary Material (Fig. S5).

**Figure 4.**
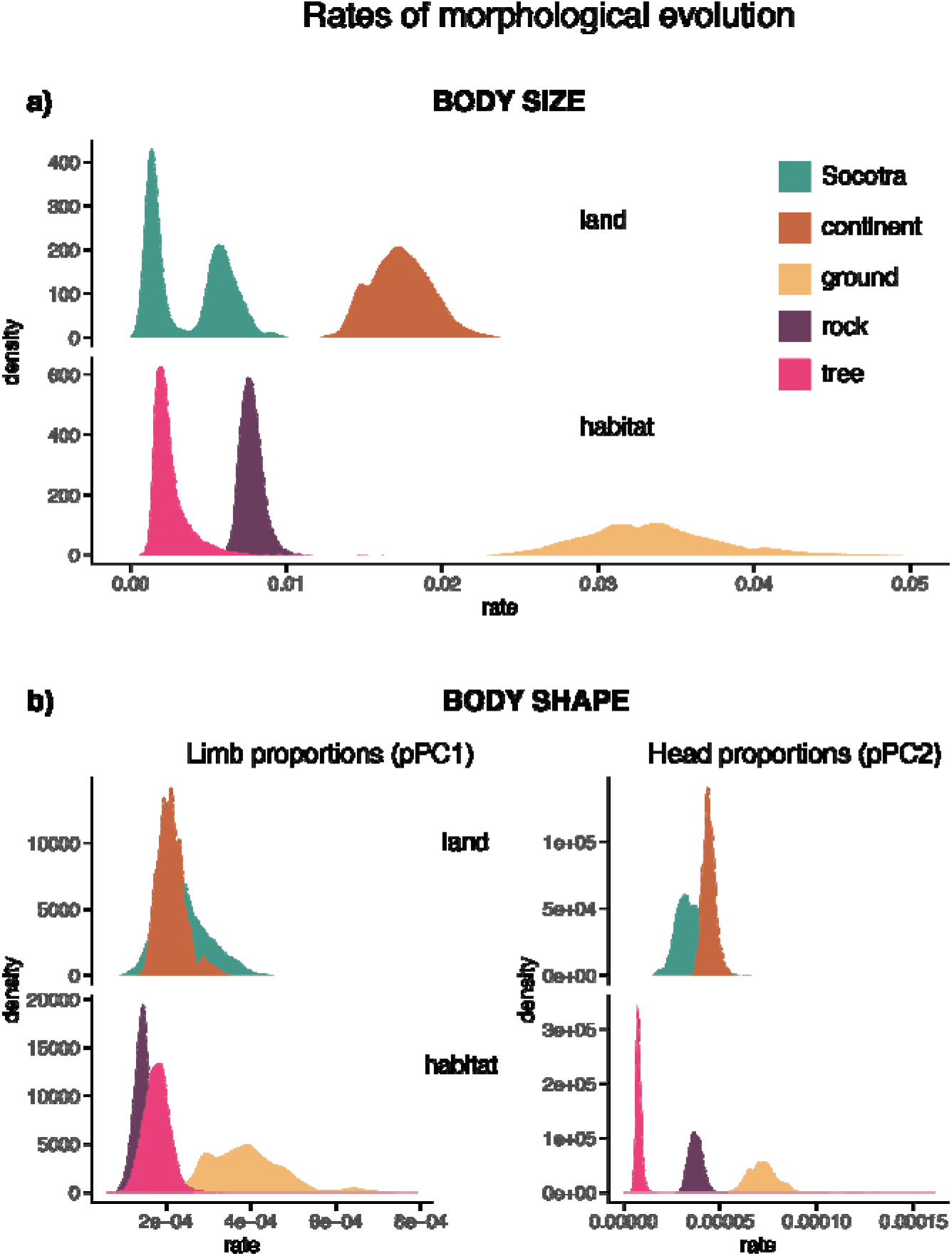
Rates of a) body size and b) body shape evolution in the genus *Pristurus*, extracted from multi-rate Brownian motion models fitted on a total of 10,000 character maps (100 stochastic character maps on 100 posterior trees) of Socotra vs. continent and habitat use (ground, rock, and tree).

## DISCUSSION

The present study represents the first comprehensive comparative work on the genus *Pristurus*, including undescribed diversity and extensive morphological (size and shape) and ecological data. We tested the relative roles of the colonisation of the Socotra Archipelago and the ecological specialisation in the evolution of the phenotypic diversity observed within the genus. We did not find evidence for the existence of an island effect in this radiation of geckos, since Socotran species do not present a notably different morphology, higher disparity, nor increased rates of morphological evolution, relative to species in the continent. On the contrary, ecological specialisation emerges as a determinant factor in generating morphological diversity, with the colonisation of ground habitats as the main driver of phenotypic divergence. Our results reveal a complex scenario in which different morphological traits interact with ecological characteristics of the species in different ways, suggesting a differential relevance of body size and shape proportions for the adaptation to specific habitats.

The tendency of island taxa to diverge in morphology compared to their continental relatives is a general pattern in terrestrial vertebrates, especially concerning body size [43,44]. In fact, recent studies on Afro-Arabian geckos colonising the Socotra Archipelago found support for this island effect, particularly in the genus *Hemidactylus* [28,29]. Nevertheless, preliminary results on *Pristurus* geckos failed to find this phenomenon in this genus [28]. Here we corroborated and extended those preliminary results, incorporating the most complete phylogenetic and morphological sampling within *Pristurus* up to date. We did not find the predicted effects of thecolonisation of the Socotra Archipelago in phenotypic diversification. Even though one of the Socotran clades (the one including *P. insignis, P. insignoides* and *P*. sp. 12) has effectively undergone an increase in body size, we did not find significant differences in body size between the first nodes of each colonisation event and their direct ancestral nodes (Fig. S3). There is also no apparent divergence in body shape, with limb and head proportions of Socotran species being completely encompassed into the morphospace of continental species (Fig. 1a). Moreover, neither size nor shape disparities observed in the Socotra Archipelago are higher than those of the continental species. Finally, our analyses failed to find another expected outcome of island colonisation, as is the increase in rates of phenotypic evolution. Consistent with our results of body size divergence, in which one of the Socotran clades is shown to have undergone a body size increase (Fig. S3), we found heterogeneity in the rates of body size evolution, with some species evolving faster than others producing a bimodal distribution (Fig. 4a). Nevertheless, species in the continent show higher rates of body size evolution than all Socotran species (Fig. 4a). For body shape, our evolutionary model fitting showed no support for differences in shape rates between continental and Socotran species (BM1 was the best-fit model; Fig. 3a). When extracting the rate values for shape from the multi-rate Brownian motion model, we found little or no difference between continental and Socotran taxa (Fig. 4b). The lack of a general island effect in *Pristurus* after the colonisation of Socotra, opposed to other similar diversifications of geckos, may indicate the existence of different ecological contexts even in the same physical settings, which would imply different ecological opportunities in the same island [11]. Namely, some life-history traits may have limited niche expansion in *Pristurus* species in the Socotra Archipelago. For instance, the fact that the ground-dwelling clade of *Pristurus* is also the only one that has adopted partially nocturnal habits (Arnold 2009) may indicate a potentially related development of these two traits. This would explain that diurnal Socotran *Pristurus* have not colonised ground habitats and therefore their morphology has not changed substantially, while some nocturnal geckos that have colonised the Socotra Archipelago, like *Hemidactylus* or *Haemodracon*, show a marked phenotypic divergence [29,45]. , This is consistent with results on global insular vertebrate communities suggesting that the prevalence of the island rule is subjected to system-specific ecological and environmental dynamics [44]. Furthermore, a recent study on the anole radiation in the Greater Antilles did not find evidence for an island effect, pointing instead to ecological opportunity and key innovations as the drivers of the adaptive radiation [46]. In *Pristurus*, one of the clades colonising the Socotra Archipelago (*P. insignis, P. insignoides* and *P*. sp. 12) seems to have undergone to a certain extent an increase in body size coupled with increased evolutionary rates compared with the other Socotran *Pristurus*. This could be indicative of some kind of ecological release derived from the island colonisation but, in any case, the degree of evolutionary change of this Socotran clade is not comparable to that observed with the colonisation of ground habitats.

Following that reasoning, ecological specialisation gives us a much more nuanced insight on *Pristurus* phenotypic evolution. The relationship between habitat use and morphological traits is well recognized in lizards [3,47,48]. In fact, preceding observations on *Pristurus* geckos suggested that many morphological changes might be functionally associated with shifts in ecology and behavior [30]. Our results are consistent with this notion and provide strong evidence that novel ecological opportunities produced high levels of phenotypic disparity associated with increased rates of trait evolution in some forms of *Pristurus*, particularly the species exploiting ground habitats. Even though ground-dwelling species do not show an extremely divergent body shape relative to species inhabiting other habitats (i.e., rocky and arboreal habitats; Fig. 1a), they do comprise the largest and the smallest sizes of the genus (Fig. 1b) and show some extreme values of limb and head proportions (pPC1 and pPC2 respectively; Fig. 1a), as well as they occupy a very large portion of the genus’ entire morphospace (Fig. 1a). This extreme variability within ground-dwelling species is reflected in our disparity results. While ground species do not present higher disparity in limb dimensions (pPC1; ratio ground/no-ground < 1), they have more than twice as much disparity as all the rest of the species in body size and head proportions, with these ratios being highly significant compared to the null model generated from simulations of character evolution (Fig. 2). Furthermore, we found increased rates of body size evolution in ground-dwelling species, followed first by rocky and last by arboreal habitats (Fig. 4a). Although rate heterogeneity across habitat categories was not the best-fit model for body shape (Fig. 3b), the rate values extracted from the multi-rate Brownian motion model show a similar pattern, especially for head proportions, with highest rates in ground-dwelling species (Fig. 4b). Taken together, these results point to the existence of a morphological response to the ecological context, especially in body size. This is consistent with the idea that the main driver of morphological divergence, even in an island colonisation event, is habitat diversity [49,50]. If habitat heterogeneity in the Socotra Archipelago is lower than in continental Africa and Arabia (e.g., no particularly large gravel plains in Socotra), or if the access to those habitats is limited for *Pristurus* geckos in Socotra and not in the continent (e.g., due to the interplay of other ecological and historical factors), phenotypic evolution after island colonisation would not be as extreme as expected under the island effect framework. This could imply a tight relationship between morphology and structural habitat, a pattern observed in other Arabian geckos such as *Ptyodactylus* or *Asaccus*, where niche conservatism is associated with a very conserved morphology [17,51–54]. This would be further supported by the fact that within ground habitats, species show a clear morphological segregation between ‘hard’ and ‘soft’ substrates, suggesting a particularly conspicuous specialisation to the former (large bodies, long limbs and large heads in the hard-ground species: *P. carteri, P. collaris, P. ornithocephalus*; Fig. S4). In fact, these hard-ground species that have developed partially nocturnal habits have evolved into rough ecological analogues of some small diurnal ground-dwelling desert iguanians (e.g., the rock agamas of the genus *Pseudotrapelus*, the toad-headed agamas of the genus *Phrynocephalus*, and the yellow-spotted agama of the genus *Trapelus*), with the long limbs favouring the movement in the newly colonised grounds and with large heads able to accommodate large eyes advantageous in those open habitats. Alternatively, the lack of an island effect in Socotran *Pristurus* might be explained by climatic divergences replacing ecomorphological differentiation, or by a low morphological evolvability [28].

Another interesting result is the morphological affinities of arboreal species, especially in body size and head proportions, where they present intermediate values within a very restricted range (Fig. 1). Consistently, we found notably reduced evolutionary rates in body size and head proportions in these species relative to other habitats with the multi-rate models (Fig. 4). This might corroborate the idea of adaptive processes leading to a tight relationship between ecological traits and phenotype, since this scenario is expected if a specific habitat constrains the morphology towards optimum values of body size and shape [14].

Ultimately, our results provide evidence of the determinant role of habitat specialisation in phenotypic evolution. This has important implications for understanding the prevalence of the island effect in the context of differential ecological opportunity and, combined with previous results on other similar systems, shows the complex nature of the relationships between ecological mechanisms and morphology and their reliance on system-specific dynamics. More detailed ecological and morphological data (e.g., dietary habits, geometric morphometrics of head shape) might help for a deeper understanding of the evolutionary dynamics of this and other groups of arid-adapted lizards.

## ACKNOWLEDGEMENTS

We are very grateful to J. Roca, M. Metallinou, K. Tamar, J. Šmid, R. Vasconcelos, R. Sindaco, F. Amat, Ph. de Pous, L. Machado, J. Garcia-Porta, J. Els, T. Mazuch, T. Papenfuss, B. Burriel and all the people from the Environment Authority, Oman, for their help in different aspects of the work. This work was supported by grants CGL2015-70390-P (MINECO/FEDER, UE) and PGC2018-098290-B-I00 (MCIU/AEI/FEDER, UE), Spain and grant 2017-SGR-00991 from the Secretaria d’Universitats i Recerca del Departament d’Economia i Coneixement de la Generalitat de Catalunya to SC. H.T.-C. was funded by an FPI grant (BES-2016-078341) (MINECO/AEI/FSE), Spain. I.M. was funded by a predoctoral grant from the Complutense University of Madrid (CT27/16-CT28/16), Spain, and the project PGC2018-094955-A-I00 from the Spanish Ministry of Science and Innovation. M.S.-R was funded by an FPI grant (BES-2013-064248) (MINECO/AEI/FSE), Spain.

## DATA AVAILABILITY STATEMENT

Supplementary material, data and R scripts used for this study are available at the online public repository https://doi.org/10.5061/dryad.xwdbrv1f6.

