## Supplementary figures and images for "Ecological specialisation, rather than the island effect, explains morphological diversification in an ancient radiation of geckos" (from another version of this preprint: version 1, 2021-07-31)

### Fig. S1

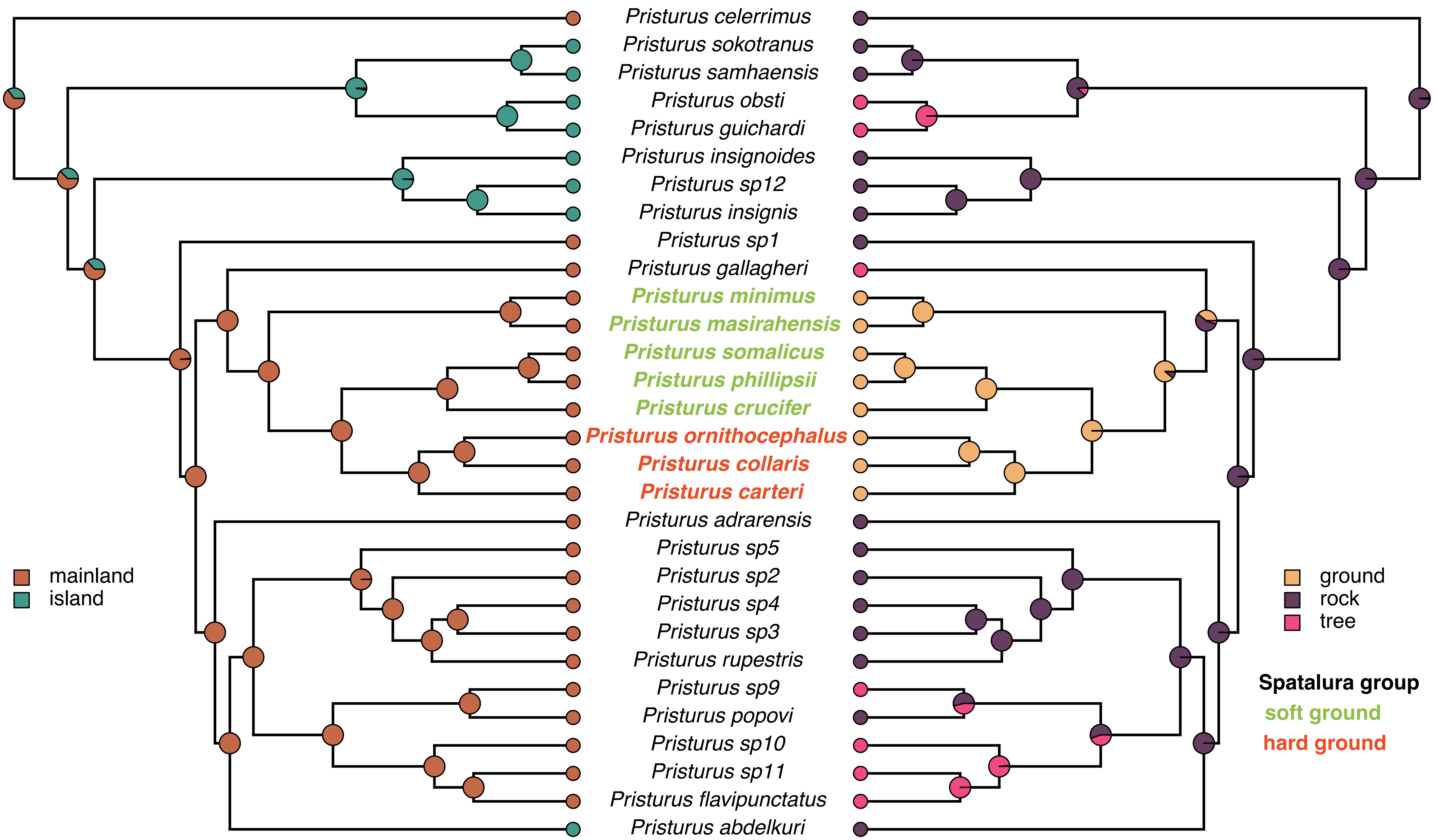

### Fig. S2

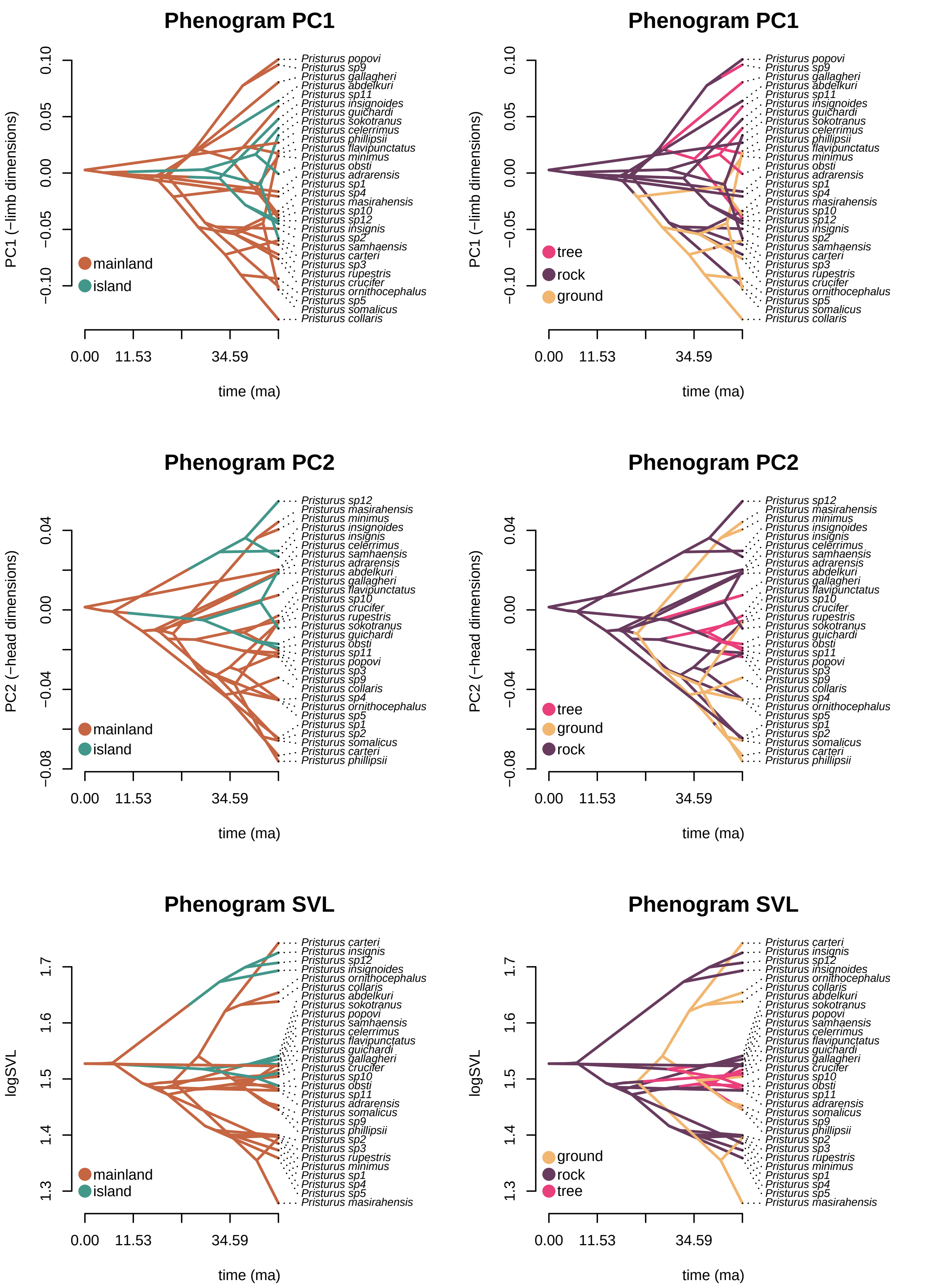

### Fig. S3

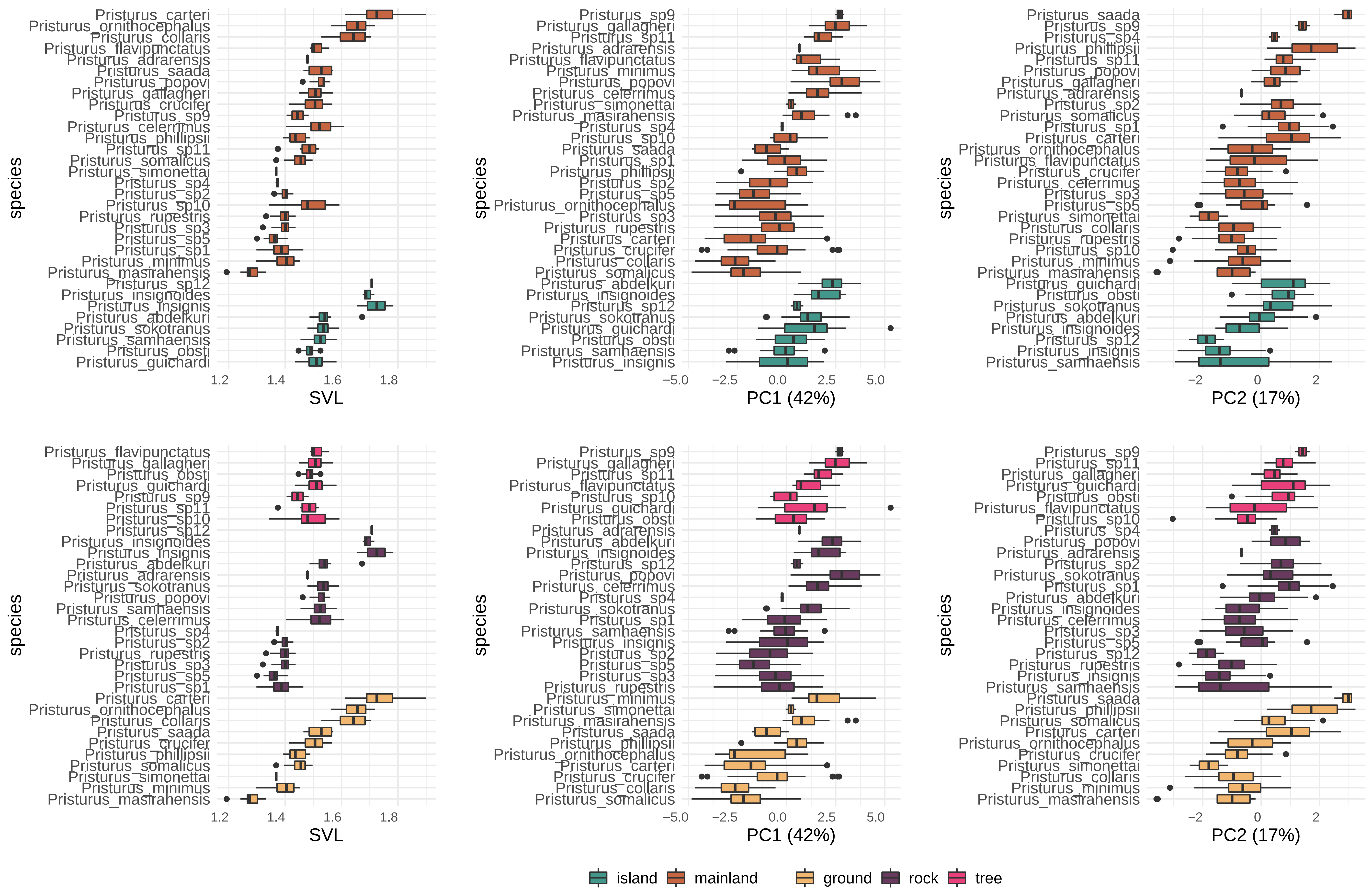

### Fig. S4

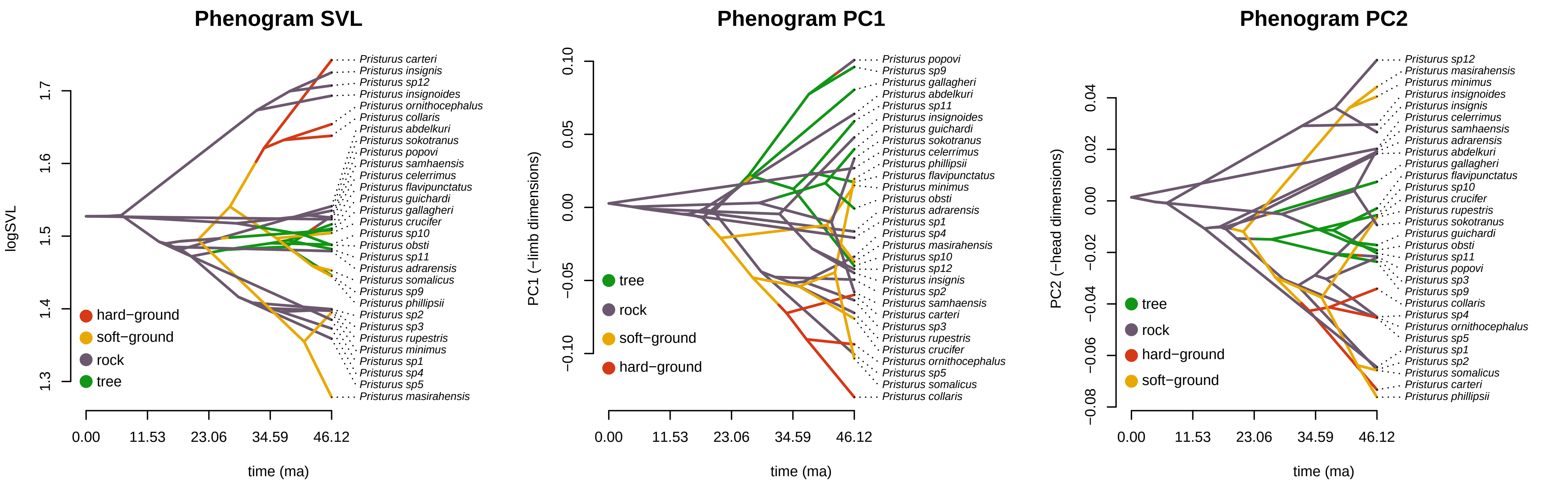

### Fig. S5

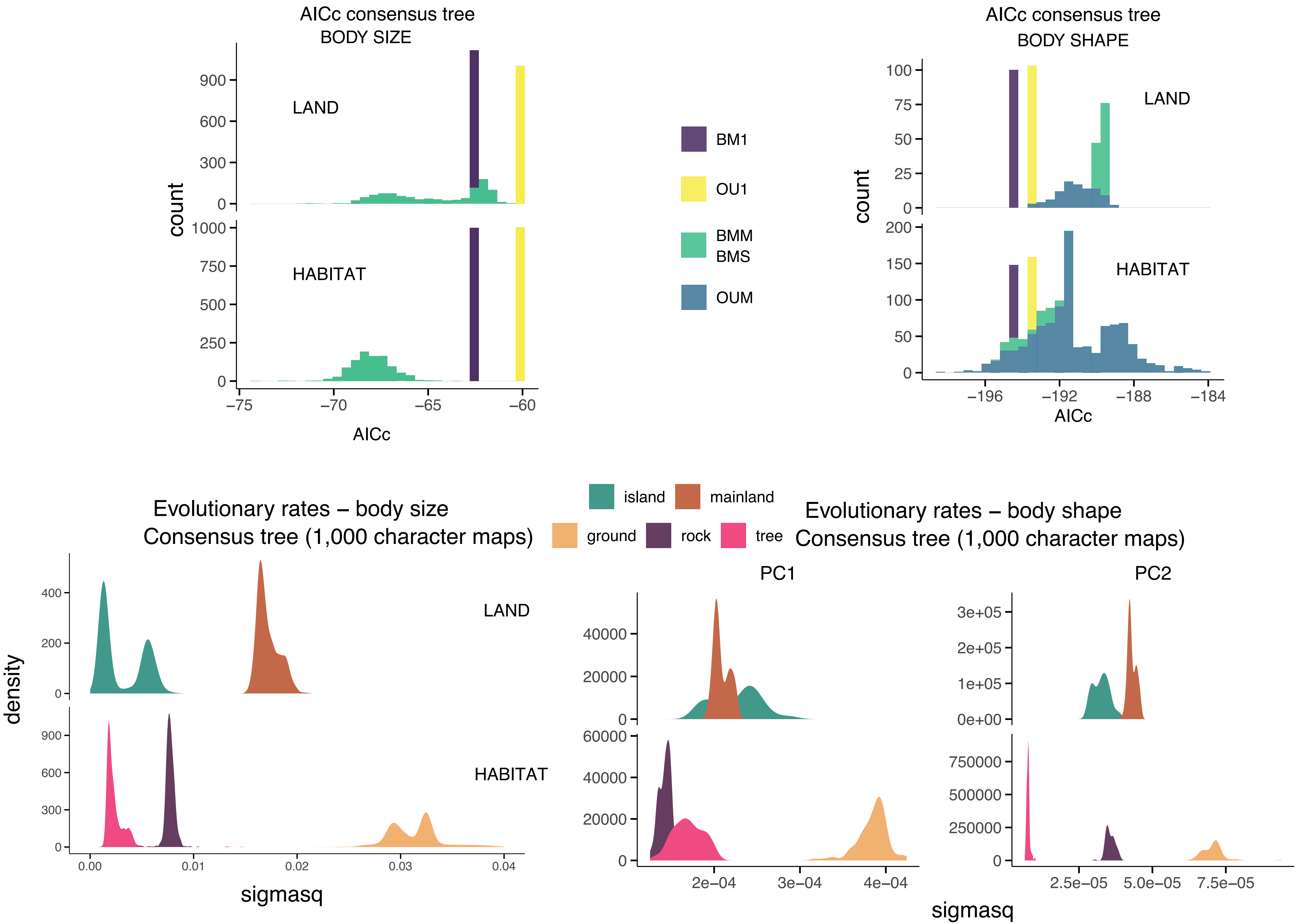

### Table S2

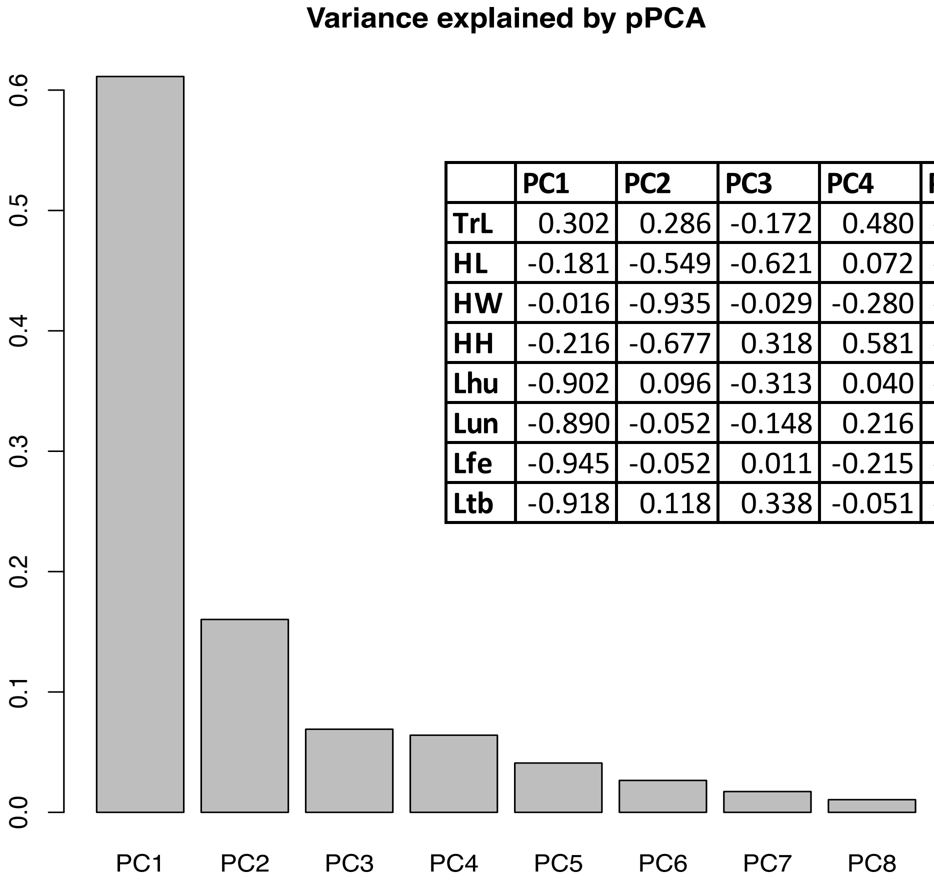


| | PC1 | PC2 | PC3 | PC4 | PC5 | PC6 | PC7 | PC8 |
|---|---|---|---|---|---|---|---|---|
| **TrL** | 0.302 | 0.286 | -0.172 | 0.480 | -0.473 | 0.550 | -0.146 | 0.139 |
| **HL** | -0.181 | -0.549 | -0.621 | 0.072 | -0.135 | -0.075 | -0.384 | -0.322 |
| **HW** | -0.016 | -0.935 | -0.029 | -0.280 | -0.028 | 0.191 | 0.097 | 0.018 |
| **HH** | -0.216 | -0.677 | 0.318 | 0.581 | -0.124 | -0.200 | 0.002 | 0.037 |
| **Lhu** | -0.902 | 0.096 | -0.313 | 0.040 | -0.201 | -0.074 | 0.176 | -0.008 |
| **Lun** | -0.890 | -0.052 | -0.148 | 0.216 | 0.351 | 0.110 | -0.005 | 0.021 |
| **Lfe** | -0.945 | -0.052 | 0.011 | -0.215 | -0.071 | -0.088 | -0.163 | 0.138 |
| **Ltb** | -0.918 | 0.118 | 0.338 | -0.051 | -0.070 | 0.100 | -0.001 | -0.107 |
